# Oral bacteriophages are maintained at high levels for months in individuals but infrequently transmitted between mothers and infants

**DOI:** 10.1101/633727

**Authors:** Clifford J. Beall, Rosalyn M. Sulyanto, Ann L. Griffen, Eugene J. Leys

## Abstract

In this work, we exploit recent advances in metagenomic assembly and bacteriophage identification to describe the phage content of saliva from 5 mother-baby pairs sampled twice 7 - 11 months apart during the first year of the babies’ lives. We identify 25 phage genomes that are comprised of one to 71 contigs, with 16 having a single contig. At the detectable level, phage were sparsely distributed with the most common one being present in 4 of the 20 samples, derived from two mothers and one baby. However, if they were present in the early time point sample from an individual, they were also present in the later sample from the same person more frequently than expected by chance. The nucleotide diversity (π) in phage from the same sample or the same person was much lower than between different individuals, indicating dominance of one strain in each person. This was different from bacterial genomes, which had higher diversity indicating the presence of multiple strains within an individual. We identify likely bacterial hosts for 16 of the 25 phage, including an apparent inovirus that is capable of integrating in the dif site of *Haemophilus* species. It appears that phage in the oral cavity are sparsely distributed, but can be maintained for months once acquired.

## INTRODUCTION

The microbiota of the oral cavity has great importance in oral health. The mechanism of dental caries involves acid production by specific bacterial populations in response to dietary sugar (1). Chronic periodontitis is likewise associated with altered bacterial communities (2, 3). Bacteriophages are likely to be an important influence on oral bacterial communities, both through predation of bacteria and aiding horizontal gene transfers. Additionally, it is possible that bacteriophages or lysins could be used as therapy for oral diseases (4).

In recent years, the availability of high throughput sequencing of the 16S rRNA gene together with other techniques has produced a wealth of information on the bacterial component of the oral microbiome (5). Less is known about oral bacteriophages, partially due to their lack of a universally conserved marker gene. However some phage that infect oral bacteria have been cultivated (4) and oral phage have also been identified in metagenomic studies (6). Most such metagenomic studies involved isolation of particles and sequencing of the particle-associated DNA, often following amplification (7–10). It has also been found that oral bacteria respond to phage with Clustered Regularly Interspaced Short Palindromic Repeat (CRISPR)-based adaptive immunity (11, 12). Recently methods have been developed to aid the identification of phage sequences from whole shotgun metagenomes without the need for particle isolation or amplification (13, 14). Paez-Espino and co-workers applied their method to whole metagenomes that were generated by the NIH Human Microbiome Project, including sequences from various human oral environments, most frequently tongue dorsum, buccal mucosa, and supragingival plaque (15).

A technique that has been developed recently to assemble bacterial genomes from metagenomes is to assemble metagenomes from many samples and then cluster contigs based on sequence coverage per sample and nucleotide kmer frequencies (16). We set out to combine the contig clustering with phage identification to identify phage genomes from whole shotgun metagenomes from the oral cavity. In the present work, we apply this approach to a data set of whole metagenomic DNA from saliva of 5 mothers and their 5 babies during the first year of the babies’ lives.

## METHODS

### Sample Collection

DNA samples were part of a larger set collected for 16S rRNA gene amplicon sequencing (Sulyanto *et al.*, in preparation). There were 20 samples, 2 per individual from 5 mother child pairs taken during the first year of the child’s life. Demographic information on the subjects is shown in Table 1. For each person we used one sample taken between 0-3 months after the birth and one sample taken 10-12 months after. Informed consent was obtained and the study was approved by The Ohio State University IRB. Saliva samples were collected from infants by saturating a flocked swab (Copan Diagnostics, www.copanusa.com) for 30 seconds and from mothers by expectoration of unstimulated saliva. Samples were stored in ATL lysis buffer and frozen until processing.

**Table 1).** Demographic Characteristics of Families.

| Family | Mother's Age | Delivery | Race | Child Gender | Child sampling months | Mother sampling months |
| --- | --- | --- | --- | --- | --- | --- |
| 1 | 28 | Vaginal | African-American | Female | 2,11 | 2,10 |
| 2 | 28 | Vaginal | Caucasian | Female | 3,10 | 3,12 |
| 4 | 27 | C-section | African-American | Male | 3,11 | 3,11 |
| 6 | 25 | Vaginal | Caucasian | Female | 3,10 | 0,11 |
| 7 | 44 | C-section | African-American | Female | 2,11 | 2,11 |

### DNA and library preparation

DNA was isolated with QIAamp DNA mini kits (QIAgen, www.qiagen.com) using the included protocol augmented by the inclusion of a bead-beating step to increase bacterial cell lysis. 100 ng of DNA in a volume of 50 µl was subjected to fragmentation in a Covaris S2 instrument with Intensity 5, Duty cycle 10%, Cycles per burst 200 and Treatment time 50s. 40 ng of the fragmented DNA was used to make Illumina sequencing libraries with the NEB Next DNA library kit for Illumina (New England Biolabs, www.neb.com). For 3 samples (family 1 mother 10 month, family 2 baby 3 month, and family 2 baby 10 month) we performed enrichment with the NEB Next Microbiome Enrichment Kit (New England Biolabs, www.neb.com) and sequenced both the pre-enrichment and post-enrichment samples. For one sample (family 1 mother 2 month), we sequenced only an enriched sample. Comparisons of the pre- and post-enrichment samples with Metaphlan2 (17) showed only minor differences in the bacterial content of the two samples so we pooled sequence data from the three repeated samples and used only the enriched data for the one sample.

We sequenced the pooled libraries in one lane of the HiSeq 4000 (Illumina, www.illumina.com) with 150 PE chemistry.

### Sequence filtering and assembly

The sequencing reads were trimmed to remove adapter sequences and low quality regions with Trimmomatic v. 0.32 (18), using the parameters ILLUMINACLIP:<TruSeq3-PE- 2.fa>:2:30:10:1:true and MINLEN:70. Human reads were removed by mapping against the human genome (human_g1k_v37.fasta from ftp://ftp.ncbi.nlm.nih.gov/1000genomes/ftp/technical/reference/) with BWA-MEM 0.7.8 (19), and processing with FilterSamReads and SamToFastq from Picard tools 1.112 (github.com/broadinstitute/picard). The reads from all samples were co-assembled using MEGAHIT v1.0.6-3-gfb1e59b (20). The assembly was examined with metaquast (21). The human-depleted sequences are deposited in the NCBI SRA associated with BioProject accession PRJNA448135.

### Clustering of contigs and identification of phage-encoding contigs

We processed the contigs through the CONCOCT clustering pipeline (16), which incorporates the following steps: (1) Co-assembly of all samples using MEGAHIT (20); (2) Discarding contigs shorter than 1 kb; (3) Fragmenting contigs larger than 20 kb into pieces of 10 kb; (4) Mapping of reads to contigs using bwa mem (19); (5) Calculation of coverage per sample for each contig; (6) Clustering of contigs into bins based on tetranucleotide frequencies and coverage per sample; and (7) Assigning taxonomy by predicting encoded protein sequences and searching against the NCBI nr non-redundant protein database. A product of the CONCOCT pipeline was a table of average coverage per sample for all assembled contigs, which we were able to use to drive decisions grouping contigs into clusters representing likely genomes.

We identified potential phage-derived contigs with VirSorter (13), and by examining the taxonomy of encoded proteins (performed with DIAMOND v. 0.8.14 (22)) and MEGAN v. 6.5.10 (23) to find bacteriophage-related genes. We further relied on the coverage per sample patterns to link phage contigs as we found that identified phage were nearly always > 2x coverage in a very small number of samples (between 1 - 4 of the 20) and < 0.1x coverage in the others. We noticed that many contigs found by VirSorter were ones that had been fragmented because they were originally over 20 kbp. We included these after checking that the coverage per sample patterns were consistent for all the fragmented pieces. We included a contig that was identified as viral and circular by VirSorter, and was 8218 bp in length (Oral phage 4), theorizing that it might be a short circular phage genome. If long contigs contained phage-related genes such as terminase or capsid proteins by blastx, we included them. We next examined CONCOCT bins that were enriched in potential phage contigs as identified by VirSorter, and included contigs from these bins if their coverage per sample levels were consistent. We split up these bins or discarded contigs if they appeared to show more than one coverage pattern. Finally we found contigs through the DIAMOND/MEGAN analysis that were related to cultivated phages. The coverage per sample for contigs in these groups were manually inspected and the contig sets were subdivided if multiple coverage patterns were found. The phage contigs are deposited at IMG/M with the genome ID 3300019854, and Supplemental Table 1 contains a mapping of the scaffold IDs in IMG/M to oral phage numbers used here and contig numbers assigned by MEGAHIT.

### Analysis of single nucleotide polymorphisms

For phage that were found in more than one sample at greater than 1 x average coverage, we analyzed the frequency of single nucleotide polymorphisms within and between samples. A step of the CONCOCT pipeline had been to align all the reads to all assembled contigs using bwa mem v. 0.7.12-r1039 (19). We used samtools v. 0.1.19 (24) to subset bam files for each sample and phage combination that exceeded the coverage threshold and generate pileup files from them. We then compared the pileups between samples at positions where each had been sequenced at least four times, and counted nucleotide differences if the position was sequenced as one base four times in one sample and a different base at least four times in the second sample. Because there is substantial overlap between the forward and reverse reads in our data (average insert size from alignment ∼140 bp), this criteria ensures the base is read in at least two independent paired end reads. We used the python script *pileup-analyze.py* to count nucleotides that were the same between samples or ones that were different.

### Nucleotide diversity and comparison of phage with bacterial genomes

To perform comparisons between bacterial genomes and phages, we selected eight contigs that were representative of four bacterial genomes, with 2 contigs corresponding to each genome from a set of abundant salivary bacteria: *Rothia mucilaginosa*, *Prevotella melaninogenica*, *Neisseria mucosa* (closely related to *Neisseria sicca* and *Morococcus cerebrosus*), and *Veillonella atypica*. We chose contigs that were over 2 kbp in length and had megablast alignments to reference genomes with 100% query coverage and over 93% identity. We then used a combination of samtools (24), BioPython (25), and command line utilities to generate bam files containing mapped reads from all samples that had over 1 fold average coverage for those contigs. We used freeBayes v1.1.0-54-g49413aa (26) to predict variants in frequency-based mode, with filters of 4 observations and 1% of total observations (parameter settings: ‘--min-alternate-fraction 0.01 --min-alternate-count 4 --pooled-continuous --haplotype-length 0’). We calculated the nucleotide diversity π as described (27), within each sample and between pairs of samples with greater than 1 fold average coverage using a python script calculate_pis.py.

### Phage annotation and taxonomy

Phage genes and proteins were predicted on the PhAnToMe annotation server (www.phantome.org). Functional annotation was carried out on the IMG/MER server (28)(available at https://img.jgi.doe.gov under IMG genome ID 3300019854).

Oral phage taxonomic relationships were analyzed with the vConTACT program (29), part of the iVirus suite on CyVerse. Briefly, the encoded proteins of the oral phage together with with the encoded proteins of all prokaryotic phage in RefSeq are used in an all versus all blastp search, the results are used to define protein clusters, and the degree of sharing of members of the protein clusters is used to define a network of relationships between phage genomes.

### Host inference by tRNA

We identified tRNA genes within the phage contigs with tRNAscan-SE 1.3.1 (30) and used the predicted sequences in blastn (31) searches of the nt and RefSeq genome databases.

### Host inference by CRISPR spacer analysis

We used two methods to attempt to identify bacterial hosts through CRISPR spacer matches with the phage genomes. The first was to search the CRISPR spacer database derived from bacterial genomes at the IMG/VR site (32) with the phage contigs. The implementation used the web interface accessed through a python script, *img-crispr-blast.py*, that invokes the Selenium web scraping library to control the Firefox browser. In the second method, we assembled CRISPR repeats from the sequence reads used in the current study using crass v. 0.3.12 (33). We then performed a blastn search with the predicted spacers as queries, the phage genomes as database, and settings ‘-task blastn-short -perc_identity 95 -qcov_hsp_perc 95 -evalue 0.1’. Because crass only assembles the CRISPRs but not flanking sequences we extracted the read ids for reads that contained the matched spacer. We then determined which MEGAHIT assembled contig they mapped to, and examined the Diamond search results of proteins encoded by the contig to assess the taxonomy.

### Host analysis - prophages

We searched for related prophages in sequenced bacterial genomes by BLAST. The refseq_genomes database was downloaded from the NCBI ftp site in December 2017 and the genetic identifier numbers (gi numbers) for bacterial sequences were accessed by querying the NCBI Entrez nucleotide site with the search term taxid2[ORGN]. We used the *blastdb_aliastool* program to create a binary form of the gi list and used blast 2.6.0+ to align the phage contigs against the database with the modifiers *-gilist <gi list file> -task blastn -evalue .00001*. Phage genomes where over 50% of the genome aligned with the bacterial genomes at over 70% nucleotide identity were considered as possible matches. The search also identified two phage that had highly similar regions (> 90% identity) to bacterial isolate genomes confined to the ends of contigs, suggesting that the metagenomic assembled contig might represent a prophage sequence.

### Clustering with metagenomic phage contigs from the “Earth’s virome” study

To determine the relationship of phage identified in this study to those found in a recent work by others (15), we applied a clustering approach that the same group developed (14). Briefly, the pipeline uses blastn to align phage genomes to the previously described metagenomic phage contigs, parses the blast results and performs single linkage clustering on the results. This allows assignment of the contigs from this study to viral clusters (vc) that they found in the earlier work and describes relationships between our contigs and singleton contigs that did not previously belong to a cluster. Since some viral clusters and singletons found in the earlier study had putative bacterial hosts, it was possible to extend this host inference to the present phage contigs.

## RESULTS

### Sequence assembly and identification of phage genomes

The sequencing produced 231 million paired end reads, 68 million of which did not map to the human genome (29%). The original assembly of those non-human reads was 221 Mbp in 200,796 contigs with an N50 value of 1189 bp. After excluding contigs under 1000 bp, there were 64,294 contigs and an assembled length of 129 Mbp. To be consistent with the recommended input for the CONCOCT program, we fragmented contigs longer than 20 kb to 10 kb pieces plus the residual.

We used the contigs greater than 1000 bp as input to the CONCOCT metagenomic clustering program (16) and it assigned them to 144 clusters. CONCOCT clusters contigs with Gaussian mixture models based on nucleotide kmer frequencies and coverage per sample. One of the outputs is a table of average coverage per sample for all contigs determined by mapping reads back to contigs with bowtie2 (34). As expected, many of these clusters represented partial to nearly complete bacterial genomes, many of them known oral bacteria. This was inferred through the CONCOCT annotation step that uses the Diamond program (22) to align predicted protein sequences against the NCBI nr protein database.

We also ran VirSorter (13) on the assembled contigs. The program identified 294 contigs using its RefSeq database and 683 contigs using the extended Virome database. Nearly all of the identified contigs were predicted to be phage-only (VirSorter categories 1-3) with only 4 contigs from the RefSeq prediction and 6 from the Virome prediction being marked as potential prophage (categories 4-6).

We had fragmented contigs that were over 20 kbp in the CONCOCT analysis pipeline to avoid possible chimeras derived from different genomes, and followed the same procedure for VirSorter. However, we noticed that a large number of the contigs identified as possible phage by VirSorter were these subfragments. We also found that the coverage per sample patterns of the subfragments were consistent with each other, indicating that they were non-chimeric. We found a total of 16 such contigs. For 14 of them, we did not find additional contigs that clustered with them in coverage per sample and they were identified as potential phage. We designated these as oral phage numbers 1-3, 5-13, 17, and 19. Another contig of 8218 bp was predicted by VirSorter to be circular by end identity. The circularity was further supported by the observation that forward and reverse ends of the same paired reads mapped at the 5’ and 3’ ends of this contig. We designated this oral phage 4. Two other >20 kb VirSorter contigs clustered with additional short contigs in phages 16 and 18.

To find additional phage, we examined CONCOCT clusters that contained higher than average numbers of predicted phage contigs by VirSorter. We found five such clusters, one of which had three distinct coverage per sample patterns that we subdivided into phage 16-18. As mentioned this resulted in additional contigs added to phages 16 and 18, while 17 remained as a single contig of 120,359 bp. For the other four CONCOCT clusters, we excluded some contigs with inconsistent coverage per sample, forming phage genomes 19-22.

We lastly examined the Diamond (22) alignments to the NCBI nr protein database to find predicted proteins that were closely related to known phage. We identified eight contigs that encoded proteins similar to Actinomyces phage Av-1 (35) (which we subdivided by coverage per sample into phage genomes 14 and 15), 9 contigs similar to Streptococcus phage Cp-1 (36) (which we subdivided into phages 24 and 25), and 4 contigs related to Streptococcus phage EJ-1 (37) (phage 23). The properties of the identified phage genomes are shown in Table 2. The collection of phages show a wide range of properties, with sizes from phage 4 at 8.2 kb to phage 20 at 231 kb and GC content from phage 22 at 23% to phage 11 at 68%.

**Table 2).** Properties of identified phage genomes.

| Phage | Num. contigs | GC content | Total length (nt) |
| --- | --- | --- | --- |
| Oral Phage 1 | 1 | 42% | 23,388 |
| Oral Phage 2 | 1 | 66% | 34,698 |
| Oral Phage 3 | 1 | 39% | 55,827 |
| Oral Phage 4 | 1 | 36% | 8,218 |
| Oral Phage 5 | 1 | 45% | 24,175 |
| Oral Phage 6 | 1 | 30% | 29,334 |
| Oral Phage 7 | 1 | 54% | 34,682 |
| Oral Phage 8 | 1 | 40% | 27,620 |
| Oral Phage 9 | 1 | 36% | 31,638 |
| Oral Phage 10 | 1 | 51% | 51,764 |
| Oral Phage 11 | 1 | 68% | 34,007 |
| Oral Phage 12 | 1 | 37% | 39,582 |
| Oral Phage 13 | 1 | 35% | 31,696 |
| Oral Phage 14 | 7 | 52% | 13,538 |
| Oral Phage 15 | 1 | 56% | 19,627 |
| Oral Phage 16 | 3 | 65% | 38,336 |
| Oral Phage 17 | 1 | 56% | 120,359 |
| Oral Phage 18 | 2 | 64% | 31,247 |
| Oral Phage 19 | 1 | 38% | 43,776 |
| Oral Phage 20 | 71 | 34% | 231,334 |
| Oral Phage 21 | 3 | 27% | 14,261 |
| Oral Phage 22 | 64 | 23% | 139,760 |
| Oral Phage 23 | 4 | 37% | 29,328 |
| Oral Phage 24 | 4 | 40% | 12,831 |
| Oral Phage 25 | 5 | 39% | 17,766 |

### Distribution of phage in samples

We summarized the read mapping data as reads mapped per genome and calculated the percentage of total reads that represented each phage. The results are plotted as a heatmap in Figure 1. In analyzing the distribution, we used a threshold of > 0.01% of reads mapping to a genome to indicate presence of a phage in a sample. Two aspects of the distribution of the phages are notable. Firstly phage are sparse with each phage only present over the threshold in a few samples. The most prevalent phage is oral phage 20, which is present in 4 of the 20 samples, from one baby and two mothers, including both early and late samples from one of the two moms. Secondly, if a phage is present in the earlier of the two samples from an individual, it has a significant tendency to be present in the other sample collected 7-11 months later (Fisher’s exact test p = 0.00012). We only found one phage that was present in both a mother and her child, *i.e.* phage 20 in family 2, in the 10 month child and 12 month mother samples.

**Figure 1).**
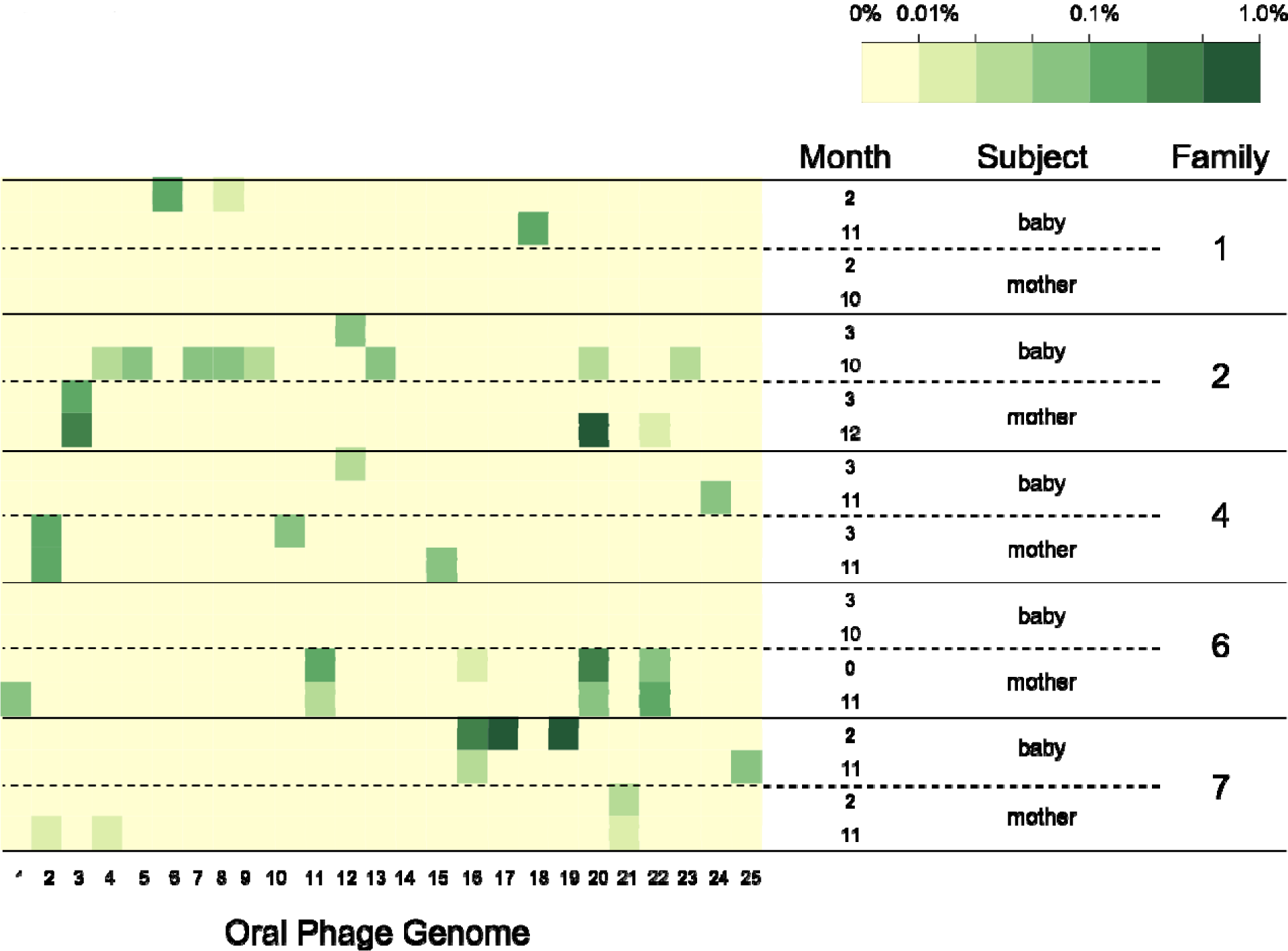
Heatmap of oral phage distribution in samples. The percentage of non-human sequence reads mapping to each phage is shown for the 20 samples. Coloring is as shown in the scale bar.

### Nucleotide diversity and single nucleotide variant (SNV) analysis

We calculated nucleotide diversity π for the phage genomes and as a comparison a set of 8 bacterial genomic contigs from four species, selected as described in methods. Nucleotide diversity is the probability that a given nucleotide is different for any two members of a population. The heuristics used to distinguish possible sequencing or PCR errors from definite variants and the method of calculating π are also described in the methods and derived from Schloissnig *et al*. (27). We calculated the diversity both within samples and between pairs of two samples, with the pairs subdivided into pairs from unrelated people, pairs from mother and baby the same family (as mentioned only one pair of samples for one phage fit this criterion), and pairs from the same person at different time points. Figure 2A shows the results of the analysis. Between unrelated people there was a broad distribution of nucleotide diversity for both phage and bacterial genomes. The mean π was 0.0055 for bacteria and 0.0048 for phage. The distribution of bacterial nucleotide diversity between family members was not significantly different from unrelated persons (mean π = 0.0058), likewise the one phage (phage 20) that was found in mother and baby had high between sample diversity. For the pairs of samples from the same person at different times, the nucleotide diversity for bacteria (mean π = 0.0029) was significantly lower than for bacteria from family members or unrelated, which agrees with similar measurements in gut (27) and skin (38) microbiomes. The diversity within samples for the bacterial contigs was lower still, though still measurable (mean π = 0.0014), indicating the presence of multiple strains for bacterial species in the oral cavity. The presence of multiple strains of oral bacterial species in single subjects has been observed in culture (39) and single cell sequencing (40, 41) experiments. Conversely the phage genomes had very low nucleotide diversity in samples from the same person, even when compared over time (mean π = 0.00034 from the same sample, = 0.00054 from different samples). Because of these nucleotide diversity patterns, in Figure 2B we treated the phage samples as single strains and analyzed the differences as percent nucleotide difference as detected by consistent variants compared to the reference assembly (which was chimeric by SNV patterns). In Figure 2B, we note that there are differences between phage species; phage 4 and phage 12 show very low divergence between subjects, while others show > 1% nucleotide difference. This includes the phage 20 mother and baby samples from family 2, which have 1.75% difference. All the within subject differences are very small, including the phage 20 samples from mother 6, which were 0.0024% different.

**Figure 2).**
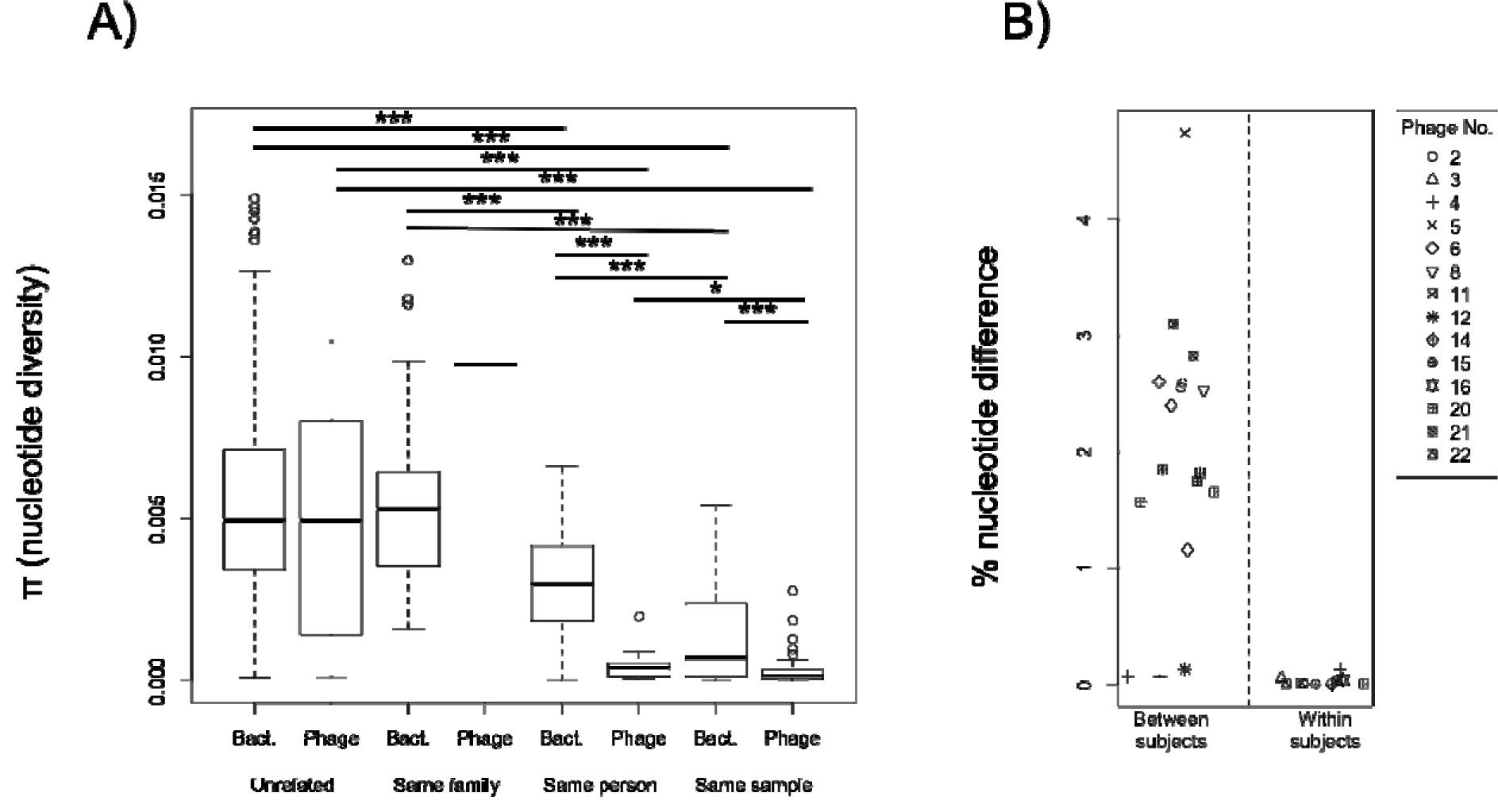
Single nucleotide variant analysis. A) Box and whisker plots of nucleotide diversity within samples (same sample) or between samples (others). The bars at top indicate significant *post hoc* Mann Whitney tests between comparison types (* p < 0.05, *** p < 0.001) B) A plot of percent nucleotide difference for phage that were found in more than one sample, based on the assumption that each sample represents a single phage strain

### Phage Taxonomy by vConTACT

We used vConTACT (29) to examine the relationship of the oral phages to cultured phages present in the NCBI RefSeq database. The program uses an all versus all protein BLAST to compare proteins and assign them to protein clusters, then determines the relationship of phage genomes by the sharing of those protein clusters. The program generates a network diagram of phage relationships based on protein clusters including the oral phage and all known phage in the RefSeq sequence database, which is presented in Figure 3. Twenty four of the twenty five oral phages had a connection to at least one RefSeq phage, the exception being oral phage 17. A group of eleven oral phages (1, 5, 6, 7, 8, 9, 12, 13, 18, 19, and 23) were well-connected to a large and interconnected group of the RefSeq phages. Phage 16 was only connected to Oral Phage 18 and a single other phage, *Arthrobacter* phage vB_ArS-ArV2 (42). This phage is part of a group of related phages that mainly infect Actinobacteria, including many infecting *Mycobacterium* and *Propionibacterium*. Another member of this cluster, *Mycobacterium* phage DS6A (43) connected to Oral Phage 18. Two other phage, Oral Phages 2 and 11, connected to a different member of this cluster, *Mycobacterium* phage PegLeg (44). However, for these four oral phages, the similarity as quantified by vConTACT between the two oral phages is stronger than the similarity between the oral phages and RefSeq phages.

**Figure 3).**
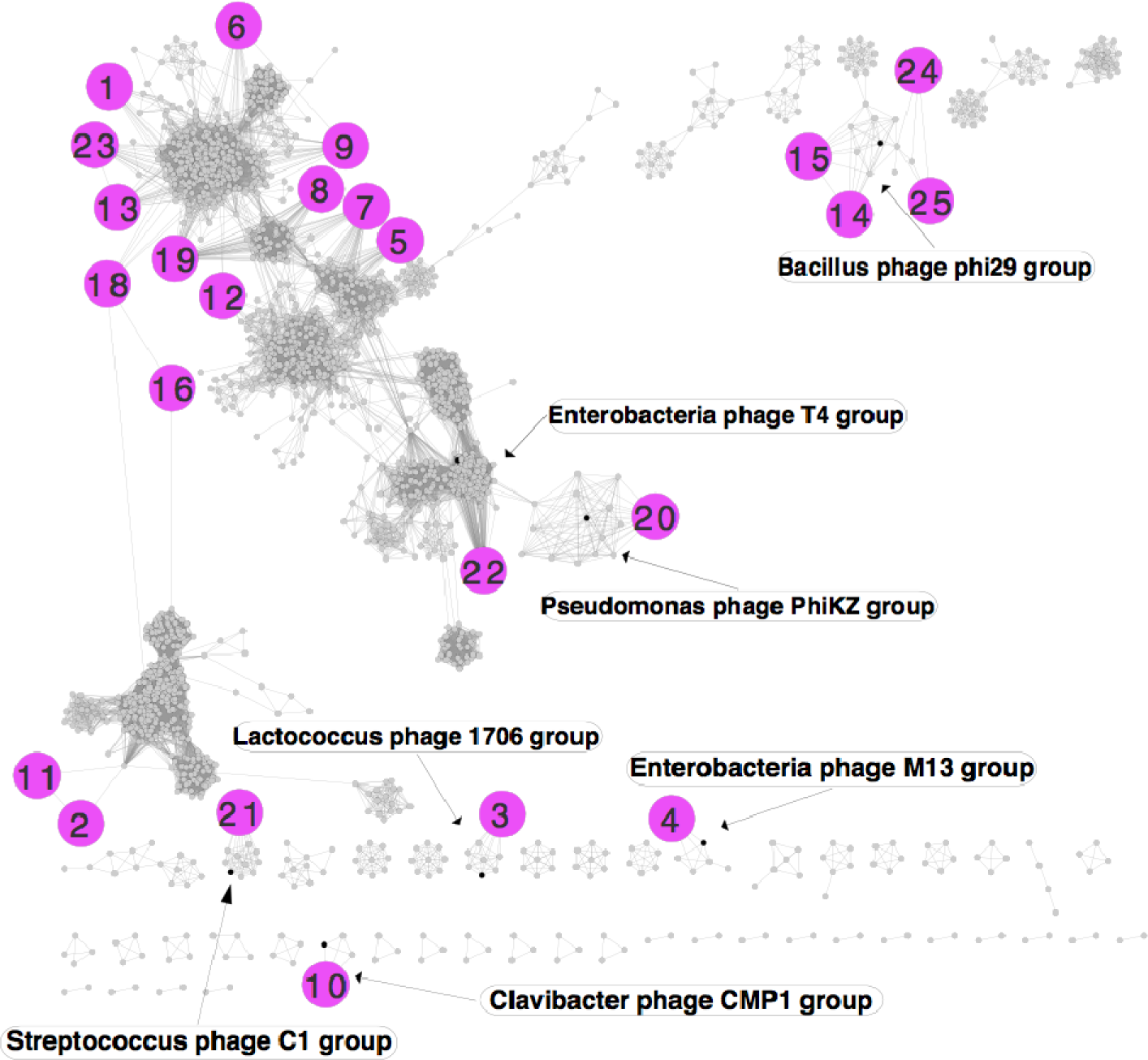
vConTACT network plot of oral phage taxonomic relationships to all RefSeq phage. The oral phage are represented by large magenta circles and have been positioned so that their connections can be seen clearly, while RefSeq phage are shown by smaller circles. RefSeq phage that characterize clusters are named and the nodes were colored black, while other RefSeq phage nodes are gray.

A group of four oral phages, numbers 14, 15, 24, and 25, are related to a cluster of phage including *Bacillus* phage phi29 (45). Of the cluster, oral phages 14 and 15 are most closely related to *Actinomyces* phage Av-1 (35), while oral phages 24 and 25 are most closely related to *Streptococcus* phage Cp-1 (46), agreeing with similarities we noted when originally identifying the phage contigs. Oral phage 20, that by genome size (> 200 kb) is considered a jumbo phage, is related to a group of jumbo phage (47) that includes *Pseudomonas* phage phiKZ. A number of phage in this group are about equally similar to oral phage 20. Oral Phage 22 is similar to a group of phage that includes the well-studied *Enterobacteria* phage T4 but has highest similarity to *Campylobacter* phage Cpt10 (48). Oral phages 3, 10, and 21 are similar to smaller clusters including less well-studied phage.

Finally, Oral Phage 4 is the only genome that shows similarity outside of the Siphoviridae (tailed phage) family. It shows similarities with members of the Inoviridae family of filamentous single stranded DNA phage, including the well known Enterobacteria phage M13.

### Phage Host Inference

The potential bacterial hosts of oral phage were inferred by four approaches: examining phage-encoded tRNA genes for potential bacterial sources, searching for CRISPR spacers that match the oral phage genomes, finding related prophage sequences in bacterial isolate genomes, and by association with previously characterized viral clusters. Table 3 summarizes the results.

**Table 3).** Host inference results.

| Oral Phage | Host by tRNA | Host by IMG/VR CRISPR | Host by crass CRISPR | Host by prophage similarity | Host by viral cluster |
| --- | --- | --- | --- | --- | --- |
| 1 |  |  |  | *Oribacterium* |  |
| 2 |  |  |  |  |  |
| 3 |  |  |  |  | *Streptococcus* |
| 4 |  |  |  | *Haemophilus* | *Haemophilus*<sup>4</sup> |
| 5 |  |  |  | *Haemophilus* |  |
| 6 |  |  | <sup>2</sup> | *Gemella* |  |
| 7 |  | Neisseriaceae | Neisseriaceae | *Neisseria* | *Neisseria* |
| 8 | Pasteurellaceae | *Haemophilus* | <sup>2</sup> | *Haemophilus* | *Haemophilus* |
| 9 |  |  | <sup>2</sup> |  |  |
| 10 | <sup>1</sup> |  | <sup>2</sup> |  |  |
| 11 |  |  | <sup>2</sup> |  |  |
| 12 |  |  |  | *Veillonella*<sup>3</sup> |  |
| 13 |  |  | <sup>2</sup> | *Streptococcus* |  |
| 14 |  |  |  |  |  |
| 15 |  |  |  |  |  |
| 16 |  | *Actinomyces* | <sup>2</sup> | *Actinomyces* | *Actinomyces* |
| 17 | <sup>1</sup> |  |  |  |  |
| 18 |  | *Actinomyces* | <sup>2</sup> |  | Actinomycetaceae |
| 19 |  | Lachnospiraceae |  | *Lachnoanaerobaculum*<sup>3</sup> |  |
| 20 | <sup>1</sup> |  |  |  |  |
| 21 |  | *Streptococcus* |  |  | *Streptococcus* |
|  |  | *sanguinis* |  |  | *sanguinis* |
| 22 |  |  |  |  |  |
| 23 | <sup>1</sup> |  | *Streptococcus* | *Streptococcus* |  |
| 24 |  | *Streptococcus* | *Streptococcus* |  | *Streptococcus mitis*<br>group. |
| 25 |  | *Streptococcus*<br>*mitis* group | *Streptococcus*<br>*mitis* group. |  | *Streptococcus mitis*<br>group |

1. Found tRNA with taxonomy not deducible
2. CRISPR spacer match with taxonomy not deducible
3. Match of ends of phage contigs to genome(s)
4. Viral cluster apparently due to second prophage in contig (see text).

The tRNAscan-SE program identified 12 potential tRNA encoding genes in the phage contigs. However, only one was a high identity match to a known bacterial tRNA. This was encoded by phage 8, and was 100% identical to tRNA-Lys genes of both *Haemophilus influenzae* and *Aggregatibacter actinomycetemcomitans*. Other tRNAs that were found were less than 95% matches to either Genbank nr/nt or RefSeq genomic databases. There were four such sequences in Oral Phage 10, three in Oral Phage 17, three in Oral Phage 20, and one in Oral Phage 23. The tRNA-containing phage that did not give host information are marked with an asterisk in Table 3.

The IMG/VR web resource (32) contains a database of CRISPR spacers derived from sequenced bacterial genomes that is searchable by BLAST. The phage contigs in this study were used to search that database and 28 hits were found to 8 of the phage. As seen in Supplemental Table 2, the BLAST matches were all to oral bacterial species. Where multiple hits were found to the same phage, the bacteria involved were taxonomically related: at genus level for phages 16, 18, 24, and 25 and at family level for phages 7 and 19.

The crass CRISPR assembly program (33) assembles CRISPR repeats from metagenomic data. The CRISPRs assembled from the metagenomic reads contained 1232 spacers, 65 of which had BLAST matches to 12 of the oral phage. Since crass only assembles the repeat regions, it was necessary to track read mappings of the repeat containing reads to attempt to identify the host by flanking genes but this was only possible for four phage. For phages 7, 24, and 25, this confirmed the identity from searching IMG/VR, while phage 23 was identified as a *Streptococcus* phage.

The fourth approach used was to search the known bacterial genomes from the RefSeq database for related sequences to the oral phage. Two kinds of sequences were found by this approach. Alignments of the entire phage genomes were found for nine of the phage genomes. We considered matches if the alignments covered at least 50% of the total length of the phage genomes, and were on average at least 70% identical. As seen in Supplemental Table 3, the percent coverage for the nine varied from 57.7% to 100% and the percent identity from 72.2% to 99.95% (rounded up in the table). Particularly striking was the high degree of similarity between Oral Phage 4 and a number of *Haemophilus influenzae* and *H. parainfluenzae* genomes. The single contig that comprises Oral Phage 4 is 8218 bp and has a direct repeat of 13bp between it’s 5’ and 3’ ends, possibly indicating a circular structure in the sample. Three of the six isolate contigs are very close to the size of the Phage 4 contig, 777_HPAR is 8259 bp, 718_HINF is 8265 bp, and HMSC068C11 is 8230 bp (Supplemental Table 3), with the difference in length reflecting longer end repeats in the isolate contigs. The other three isolates have the phage regions as part of longer contigs. In each case the contig represents a phage insertion that occurs at a position corresponding to the dif site of *Haemophilus influenzae* (49), a form of prophage that has been observed for filamentous phage of the Inoviridae family (50).

Other related prophage were found for oral phages 1, 5, 6, 7, 8, 13, 16, and 23 (Supplemental Table 3). For phages 7, 8, 16, and 23 the host inference agreed with other methods while for the other phage host predictions were not available. In the case of phage 23, the nucleotide identity was the lowest, and different contigs of the phage were most highly identical to different isolate genomes, although all similarities were to *Streptococcus* strains related to *Streptococcus mitis*.

Finally oral phages 12 and 19 showed nucleic acid identities to bacterial isolates only at the ends of the metagenomic phage contigs. The regions involved are shown in Supplemental Table 4. In the case of oral phage 12, the regions at the ends represent contiguous regions from the similar genomes, indicating that the phage may have integrated in the genome. The putative integration point is within a gene that is predicted to encode a YebC/PmpR family DNA-binding transcriptional regulator. The insertion is predicted to change the amino acid sequence at the C-terminus of the protein from AIMDEEE to SILINE. The insertion point is upstream of sequences encoding the TPP riboswitch and the ThiC gene involved in thiamine biosynthesis. Very near the 5’ end of the contig for oral phage 19 there is a sequence highly similar to sequences from *Lachnoanaerobaculum* sp. ICM7 that are predicted to encode an IS110 family transposase while the 3’ end consists of a 94 bp repeated sequence from upstream of the transposase. The predicted host of this phage by CRISPR spacer analysis was Lachnospiraceae, the family containing *Lachnoanaerobaculum*.

We searched the annotations generated by the IMG-MER system to identify possible integrase related genes, including transposases and resolvases, as these might be expected in temperate phage capable of integration. The results are presented in Supplemental Table 5. Five of the nine phage that had related prophages contained integrase or related genes (5, 6, 7, 8, and 16), as did both of the phage with ends similar to isolate genomes (12 and 20). The lack of integrase for phage 4 is not surprising since many related filamentous phage utilize cellular recombination proteins XerC and XerD to catalyze integration at the dif site rather than encoding their own integrase (50). Integrases were not identified for phages 1, 13, and 23 despite the existence of (somewhat distantly) related prophages.

### Viral cluster analysis

A recent publication identified phage-encoding contigs from a variety of sources, including metagenomic sequences from oral samples generated by the Human Microbiome Project (15). In that publication the authors described clustering of viral contigs and the method for clustering was published separately (14). Contigs are clustered based on 90% average nucleotide identity over 75% overlap. This method was applied to the phage genomes found in the current study, combining them with viral clusters and singletons found in the previous study (Supplemental Table 6). Twenty of the twenty five oral phage clustered with phage contigs found in that work. Phage 24 and 25 clustered with each other and the same viral clusters and singletons. Although those authors identified >100,000 viral clusters from numerous host-associated and other habitats, all of the contigs that clustered with phage from the current study were derived from the human oral environment.

The case of phage 4 was somewhat anomalous. The contig to which it showed high identity, metagenomic contig SRS015921_WUGC_scaffold_307 is 55,406 in length and appeared to be a segment of a *Haemophilus parainfluenzae* genome, with phage 4 inserted at the dif site as described above, and an unrelated prophage inserted about 1 kb proximal to the dif site downstream of the genomic Ferritin-1 and Ferritin-2 genes, which are over 98% identical to ones from *H. parainfluenzae*. The viral cluster that was observed in the previous study appears to be due to the presence of this unrelated phage sequence, which is likely a member of the Siphoviridae as it contains conserved tail fiber protein genes. Finally host inference was also possible through the viral clustering in some cases (Table 3), and allowed the prediction of oral phage 3 as a Streptococcus phage.

## DISCUSSION

In this work, we describe a set of oral bacteriophage that were found solely by analysis of whole metagenome shotgun sequences, without isolation of viral particles or DNA amplification. This method has some advantages and disadvantages. The advantages include that it was possible to make inferences by the coverage per sample of the assembled contigs, and associate contigs that derive from the same phage. Therefore it was possible to study phage that assembled as multiple contigs. Methods using coverage per sample that have been useful in generating bacterial genomes from metagenomes (51) can be applied to phage in this way. This was particularly true for oral samples in this study because each phage had significant coverage in only a few samples and appeared to be absent in other ones. A second advantage to the approach is that filtration methods that are sometimes used to isolate viral particles often exclude jumbo phage (47), one of which was found in this study.

The depth of sequencing of this study is somewhat limited by sequencing costs, the desire to study multiple samples including longitudinal ones, and the presence of 71% human DNA in saliva. A possibility is that phages are not as sparse as they appear, but are normally present at low levels and undergo blooms to detectable levels given specific environmental conditions. Two observations seem to speak against this possibility. One is that the same phages are very often found at detectable levels in samples from the same individual 7-11 months apart. A second is that phages within an individual at high levels have low nucleotide diversity, while between individuals most phages have higher diversity with the exception of phages 12 and 4. Phage 12 was only found in the babies from families 2 and 4 and is not as highly identical to related phage contigs in IMG/VR (90-98% nucleotide identity). Phage 4 was also found to be have high identity to various *Haemophilus* isolate genomes and a metagenomic contig assembled from the Human Microbiome Project data (32).

It is also possible that the phages studied here reflect a subpopulation of oral phages that are sparsely distributed. It may be that there are more common phages that are present in more slightly divergent forms, but the presence of multiple related genomes may interfere with genome assembly into long enough contigs to observe and reliably classify as phage.

In this study although we frequently observed maintenance of phage for months with very low nucleotide divergence between samples from the same patient we failed to find any cases of phage transfer from mothers to children. In the single case of phage 20 where a very similar phage was present in a mother and baby from the same family, the strain was as different (1.75% nucleotide divergence) as many phage from unrelated subjects, suggesting that it was acquired from a different source. Other studies have shown transmission of phage from mothers to babies (52–54) so it is likely there was not a large enough sample size of phage and subjects to observe it here. Transmission does seem to be less frequent than maintenance within an individual.

A drawback of using the whole metagenome approach is that it is not possible to know for certain if the phage sequences identified are prophage inserted in bacterial genomes, some other intracellular form, or phage particles. The evidence suggests there may be some of all types. Phage 4 is an interesting case. The method that was used to create sequencing libraries involving repairing DNA ends, tailing, and ligating adapters. Such a technique would not be expected to work with single stranded DNA. Phage 4 is related to the Inoviridae family of filamentous phage where the packaged genome is a single stranded DNA circle. The lifecycle of these phages however involves an intracellular double stranded circular replicative form, and it may be this form is observed here. Some contigs from bacterial isolates and a metagenomic contig strongly suggest that an integrated form of phage 4 in the dif site of Haemophilus species is possible, though we did not directly observe it. Phages 12 and 19 had ends that were highly identical to bacterial genomes, suggesting that the contigs might be derived from prophage sequences, especially in the case of 12 where contiguous sequences from *Veillonella* genomes that contained metabolic genes were present at either end. The case for phage 19 was not quite so obvious because the sequence at one end was a repeat of part of the sequence from the other, the similarity was only with a single isolate genome, and the region involved contained a transposase gene. It is possible a transposon inserted into a phage genome and was packaged. However the species of bacteria agreed with the predicted host of the phage, indicating the fusion did not result from a misassembly.

Several approaches were used to identify the bacterial hosts for these oral phage. Altogether we were able to identify putative hosts for 16 of the 25 phage through tRNA analysis, CRISPR spacer analysis, similarity to prophages, and clustering with previously described phage (Table 3). Where it was possible to identify bacterial hosts by multiple methods, there was agreement between the methods though in some cases one method giving a higher taxonomic rank that subsumed the host prediction by the other method. This gives us confidence in the predictions.

The viral clustering indicates that this study represents a fairly small subset of possible oral phage. From the IMG/VR site (32), it is reported that they found 48,904 viral contigs and associated those with 5246 viral clusters and had remaining 7467 singleton contigs. The phage genomes from our study associated with 19 of those clusters and 31 singletons. Our oral phage also would have united some clusters and singletons from the earlier study. In all cases the phage identified here associated with human oral phage. Given that the criterion for clustering is fairly stringent, the fact that 18 of 25 of the phage in this study clustered with the IMG/VR phage indicates that collection may be fairly comprehensive (note that phage 4 may have been included serendipitously).

It would be of interest to identify phage that could be utilized in phage therapy to treat oral diseases or opportunistic infections caused by oral bacteria (4). It would require additional work to cultivate such phage but the current work could be used to develop rapid PCR tests for phage presence in samples and in some cases such as phage 4 identify bacterial strains containing prophage. We identified multiple phages that appear to infect *Streptococcus*, though none that were shown to directly target *S. mutans* or *S. sobrinus*, which have significant roles in dental caries. Also we did not find phage that obviously infect bacteria strongly associated with periodontitis such as *Porphyromonas gingivalis*, *Treponema denticola*, *Tannerella forsythia*, *Filifactor alocis*, or others (2). It might be possible to more efficiently find such phage by examining DNA isolated from supragingival plaque for caries or subgingival plaque for periodontitis. However bacteria including species from *Streptococcus*, *Haemophilus*, and *Actinomyces* can cause opportunistic infections and we discovered phage infecting each of those genera.

In conclusion, we have shown that bacteriophage can be readily discovered by analysis of whole metagenomic sequences from the oral cavity. We find that phage if present seem to have low nucleotide diversity, indicating that they are dominated by a single strain, a striking difference from oral bacteria which have higher nucleotide diversity indicating the presence of multiple strains in most cases. Phage appear to be sparsely distributed but if present are frequently maintained over periods of months.

## ACKNOWLEDGEMENTS

This work was supported by NIH/NIDCR grant DE024327. We thank Pearlly Yan for DNA sequencing, Haella Holmes and Steve Spiritoso for technical assistance, Ben Bolduc for help with vConTACT and Ann Gregory and Matthew Sullivan for useful discussions.

## SOFTWARE AND DATA SHARING

Software scripts used in the work are available as a git repository at https://code.osu.edu/beall-3/salivary-virome. Raw sequences are available at NCBI SRA under BioProject PRJNA448135. Assembled phage contigs are available at https://img.jgi.doe.gov/ under IMG genome ID 3300019854.

---

**Supplemental Table 1:** Mapping of scaffold IDs in IMG/M to oral phage numbers and MEGAHIT contigs in this study.

| IMG Scaffold | Sequence Length(bp) | GC Content | Gene Count | MetaHIT contig number | Oral Phage Number |
| --- | --- | --- | --- | --- | --- |
| 3300019854 assembled Ga0206370_1001 | 1398 | 0.51 | 2 | k99_33649 | 14 |
| 3300019854 assembled Ga0206370_1002 | 1685 | 0.54 | 2 | k99_148626 | 14 |
| 3300019854 assembled Ga0206370_1003 | 2822 | 0.54 | 3 | k99_458922 | 14 |
| 3300019854 assembled Ga0206370_1004 | 1032 | 0.52 | 2 | k99_192535 | 14 |
| 3300019854 assembled Ga0206370_1005 | 3046 | 0.54 | 3 | k99_98426 | 14 |
| 3300019854 assembled Ga0206370_1006 | 2055 | 0.51 | 5 | k99_260963 | 14 |
| 3300019854 assembled Ga0206370_1007 | 1500 | 0.46 | 4 | k99_408416 | 14 |
| 3300019854 assembled Ga0206370_1008 | 8218 | 0.36 | 13 | k99_304421 | 4 |
| 3300019854 assembled Ga0206370_1009 | 34007 | 0.68 | 44 | k99_406096 | 11 |
| 3300019854 assembled Ga0206370_1010 | 9740 | 0.36 | 14 | k99_375836 | 23 |
| 3300019854 assembled Ga0206370_1011 | 14711 | 0.37 | 22 | k99_434423 | 23 |
| 3300019854 assembled Ga0206370_1012 | 3159 | 0.43 | 7 | k99_448687 | 23 |
| 3300019854 assembled Ga0206370_1013 | 1718 | 0.43 | 5 | k99_481303 | 23 |
| 3300019854 assembled Ga0206370_1014 | 3670 | 0.63 | 5 | k99_313503 | 18 |
| 3300019854 assembled<br>Ga0206370_1015 | 27577 | 0.64 | 39 | k99_368282 | 18 |
| 3300019854 assembled<br>Ga0206370_1016 | 27620 | 0.4 | 40 | k99_347722 | 8 |
| 3300019854 assembled<br>Ga0206370_1017 | 3911 | 0.36 | 7 | k99_122647 | 25 |
| 3300019854 assembled<br>Ga0206370_1018 | 1488 | 0.32 | 6 | k99_276034 | 25 |
| 3300019854 assembled<br>Ga0206370_1019 | 2047 | 0.39 | 3 | k99_317095 | 25 |
| 3300019854 assembled<br>Ga0206370_1020 | 6909 | 0.42 | 8 | k99_74971 | 25 |
| 3300019854 assembled<br>Ga0206370_1021 | 3411 | 0.39 | 5 | k99_82015 | 25 |
| 3300019854 assembled<br>Ga0206370_1022 | 23388 | 0.42 | 37 | k99_155121 | 1 |
| 3300019854 assembled<br>Ga0206370_1023 | 34698 | 0.66 | 44 | k99_16347 | 2 |
| 3300019854 assembled<br>Ga0206370_1024 | 31638 | 0.36 | 39 | k99_373793 | 9 |
| 3300019854 assembled<br>Ga0206370_1025 | 2450 | 0.24 | 7 | k99_130132 | 21 |
| 3300019854 assembled<br>Ga0206370_1026 | 8411 | 0.28 | 8 | k99_170084 | 21 |
| 3300019854 assembled<br>Ga0206370_1027 | 3400 | 0.26 | 5 | k99_468486 | 21 |
| 3300019854 assembled<br>Ga0206370_1028 | 29334 | 0.3 | 48 | k99_326664 | 6 |
| 3300019854 assembled<br>Ga0206370_1029 | 51764 | 0.51 | 93 | k99_37810 | 10 |
| 3300019854 assembled<br>Ga0206370_1030 | 2057 | 0.39 | 3 | k99_249821 | 24 |
| 3300019854 assembled<br>Ga0206370_1031 | 6813 | 0.42 | 7 | k99_293096 | 24 |
| 3300019854 assembled<br>Ga0206370_1032 | 1177 | 0.4 | 3 | k99_352772 | 24 |
| 3300019854 assembled<br>Ga0206370_1033 | 2784 | 0.38 | 3 | k99_79926 | 24 |
| 3300019854 assembled<br>Ga0206370_1034 | 3256 | 0.34 | 4 | k99_5394 | 20 |
| 3300019854 assembled<br>Ga0206370_1035 | 1038 | 0.28 | 2 | k99_6686 | 20 |
| 3300019854 assembled<br>Ga0206370_1036 | 3801 | 0.35 | 4 | k99_7314 | 20 |
| 3300019854 assembled<br>Ga0206370_1037 | 2803 | 0.33 | 5 | k99_12116 | 20 |
| 3300019854 assembled<br>Ga0206370_1038 | 8761 | 0.34 | 8 | k99_21210 | 20 |
| 3300019854 assembled<br>Ga0206370_1039 | 1356 | 0.31 | 3 | k99_21681 | 20 |
| 3300019854 assembled<br>Ga0206370_1040 | 2618 | 0.3 | 3 | k99_23954 | 20 |
| 3300019854 assembled<br>Ga0206370_1041 | 1481 | 0.38 | 4 | k99_41363 | 20 |
| 3300019854 assembled<br>Ga0206370_1042 | 8423 | 0.35 | 7 | k99_42953 | 20 |
| 3300019854 assembled<br>Ga0206370_1043 | 3710 | 0.34 | 5 | k99_48500 | 20 |
| 3300019854 assembled<br>Ga0206370_1044 | 2793 | 0.34 | 4 | k99_56643 | 20 |
| 3300019854 assembled<br>Ga0206370_1045 | 4717 | 0.36 | 4 | k99_57911 | 20 |
| 3300019854 assembled<br>Ga0206370_1046 | 4209 | 0.34 | 3 | k99_58759 | 20 |
| 3300019854 assembled<br>Ga0206370_1047 | 1008 | 0.27 | 1 | k99_64785 | 20 |
| 3300019854 assembled<br>Ga0206370_1048 | 2846 | 0.33 | 3 | k99_65434 | 20 |
| 3300019854 assembled<br>Ga0206370_1049 | 6435 | 0.34 | 11 | k99_66351 | 20 |
| 3300019854 assembled<br>Ga0206370_1050 | 3220 | 0.33 | 2 | k99_78529 | 20 |
| 3300019854 assembled<br>Ga0206370_1051 | 6393 | 0.32 | 10 | k99_89208 | 20 |
| 3300019854 assembled<br>Ga0206370_1052 | 5602 | 0.38 | 5 | k99_89405 | 20 |
| 3300019854 assembled<br>Ga0206370_1053 | 2459 | 0.34 | 4 | k99_95873 | 20 |
| 3300019854 assembled<br>Ga0206370_1054 | 1779 | 0.35 | 1 | k99_103797 | 20 |
| 3300019854 assembled<br>Ga0206370_1055 | 3694 | 0.35 | 2 | k99_104674 | 20 |
| 3300019854 assembled<br>Ga0206370_1056 | 1248 | 0.31 | 2 | k99_104866 | 20 |
| 3300019854 assembled<br>Ga0206370_1057 | 1182 | 0.32 | 3 | k99_112847 | 20 |
| 3300019854 assembled<br>Ga0206370_1058 | 3298 | 0.32 | 4 | k99_112899 | 20 |
| 3300019854 assembled<br>Ga0206370_1059 | 4617 | 0.36 | 4 | k99_123933 | 20 |
| 3300019854 assembled<br>Ga0206370_1060 | 2046 | 0.33 | 3 | k99_124329 | 20 |
| 3300019854 assembled<br>Ga0206370_1061 | 1373 | 0.31 | 2 | k99_136916 | 20 |
| 3300019854 assembled<br>Ga0206370_1062 | 3004 | 0.34 | 3 | k99_136923 | 20 |
| 3300019854 assembled<br>Ga0206370_1063 | 1165 | 0.28 | 3 | k99_138623 | 20 |
| 3300019854 assembled<br>Ga0206370_1064 | 3052 | 0.32 | 4 | k99_139067 | 20 |
| 3300019854 assembled<br>Ga0206370_1065 | 2040 | 0.31 | 3 | k99_145556 | 20 |
| 3300019854 assembled<br>Ga0206370_1066 | 3906 | 0.32 | 3 | k99_146578 | 20 |
| 3300019854 assembled<br>Ga0206370_1067 | 4944 | 0.3 | 4 | k99_146994 | 20 |
| 3300019854 assembled<br>Ga0206370_1068 | 1910 | 0.34 | 3 | k99_161151 | 20 |
| 3300019854 assembled<br>Ga0206370_1069 | 3309 | 0.34 | 5 | k99_182064 | 20 |
| 3300019854 assembled<br>Ga0206370_1070 | 1325 | 0.3 | 4 | k99_185128 | 20 |
| 3300019854 assembled<br>Ga0206370_1071 | 3791 | 0.35 | 3 | k99_186551 | 20 |
| 3300019854 assembled<br>Ga0206370_1072 | 7169 | 0.32 | 11 | k99_194323 | 20 |
| 3300019854 assembled<br>Ga0206370_1073 | 9403 | 0.33 | 8 | k99_196319 | 20 |
| 3300019854 assembled<br>Ga0206370_1074 | 3546 | 0.34 | 5 | k99_199804 | 20 |
| 3300019854 assembled<br>Ga0206370_1075 | 2746 | 0.35 | 2 | k99_201625 | 20 |
| 3300019854 assembled<br>Ga0206370_1076 | 3024 | 0.37 | 3 | k99_216693 | 20 |
| 3300019854 assembled<br>Ga0206370_1077 | 2269 | 0.32 | 2 | k99_216870 | 20 |
| 3300019854 assembled<br>Ga0206370_1078 | 3331 | 0.35 | 4 | k99_217123 | 20 |
| 3300019854 assembled<br>Ga0206370_1079 | 1873 | 0.32 | 5 | k99_219794 | 20 |
| 3300019854 assembled<br>Ga0206370_1080 | 1457 | 0.35 | 3 | k99_219969 | 20 |
| 3300019854 assembled<br>Ga0206370_1081 | 2120 | 0.34 | 3 | k99_238270 | 20 |
| 3300019854 assembled<br>Ga0206370_1082 | 5836 | 0.37 | 4 | k99_239247 | 20 |
| 3300019854 assembled<br>Ga0206370_1083 | 5098 | 0.31 | 7 | k99_248892 | 20 |
| 3300019854 assembled<br>Ga0206370_1084 | 1308 | 0.33 | 2 | k99_249212 | 20 |
| 3300019854 assembled<br>Ga0206370_1085 | 2093 | 0.32 | 4 | k99_250679 | 20 |
| 3300019854 assembled<br>Ga0206370_1086 | 9490 | 0.35 | 5 | k99_263124 | 20 |
| 3300019854 assembled<br>Ga0206370_1087 | 3408 | 0.34 | 4 | k99_268053 | 20 |
| 3300019854 assembled<br>Ga0206370_1088 | 1831 | 0.32 | 2 | k99_274402 | 20 |
| 3300019854 assembled<br>Ga0206370_1089 | 4285 | 0.31 | 4 | k99_275561 | 20 |
| 3300019854 assembled<br>Ga0206370_1090 | 3501 | 0.34 | 6 | k99_281895 | 20 |
| 3300019854 assembled<br>Ga0206370_1091 | 1113 | 0.32 | 3 | k99_310579 | 20 |
| 3300019854 assembled<br>Ga0206370_1092 | 2388 | 0.37 | 2 | k99_322695 | 20 |
| 3300019854 assembled<br>Ga0206370_1093 | 1186 | 0.34 | 2 | k99_332721 | 20 |
| 3300019854 assembled<br>Ga0206370_1094 | 1884 | 0.37 | 2 | k99_351027 | 20 |
| 3300019854 assembled<br>Ga0206370_1095 | 4844 | 0.37 | 2 | k99_364925 | 20 |
| 3300019854 assembled<br>Ga0206370_1096 | 2873 | 0.35 | 2 | k99_421298 | 20 |
| 3300019854 assembled<br>Ga0206370_1097 | 2461 | 0.34 | 1 | k99_430226 | 20 |
| 3300019854 assembled<br>Ga0206370_1098 | 1118 | 0.34 | 2 | k99_443195 | 20 |
| 3300019854 assembled<br>Ga0206370_1099 | 1705 | 0.33 | 2 | k99_443205 | 20 |
| 3300019854 assembled<br>Ga0206370_1100 | 6027 | 0.32 | 7 | k99_447517 | 20 |
| 3300019854 assembled<br>Ga0206370_1101 | 2976 | 0.34 | 7 | k99_447858 | 20 |
| 3300019854 assembled<br>Ga0206370_1102 | 1011 | 0.29 | 1 | k99_448160 | 20 |
| 3300019854 assembled<br>Ga0206370_1103 | 1144 | 0.3 | 3 | k99_451094 | 20 |
| 3300019854 assembled<br>Ga0206370_1104 | 2204 | 0.35 | 3 | k99_475083 | 20 |
| 3300019854 assembled<br>Ga0206370_1105 | 34682 | 0.54 | 59 | k99_345784 | 7 |
| 3300019854 assembled<br>Ga0206370_1106 | 19627 | 0.56 | 26 | k99_203517 | 15 |
| 3300019854 assembled<br>Ga0206370_1107 | 9193 | 0.65 | 16 | k99_66521 | 16 |
| 3300019854 assembled<br>Ga0206370_1108 | 20238 | 0.65 | 26 | k99_217445 | 16 |
| 3300019854 assembled<br>Ga0206370_1109 | 8905 | 0.65 | 20 | k99_283470 | 16 |
| 3300019854 assembled<br>Ga0206370_1110 | 39582 | 0.37 | 45 | k99_417291 | 12 |
| 3300019854 assembled<br>Ga0206370_1111 | 1710 | 0.22 | 3 | k99_1529 | 22 |
| 3300019854 assembled<br>Ga0206370_1112 | 3856 | 0.2 | 8 | k99_5647 | 22 |
| 3300019854 assembled<br>Ga0206370_1113 | 2247 | 0.22 | 4 | k99_11194 | 22 |
| 3300019854 assembled<br>Ga0206370_1114 | 2467 | 0.25 | 4 | k99_11326 | 22 |
| 3300019854 assembled<br>Ga0206370_1115 | 1575 | 0.23 | 3 | k99_36403 | 22 |
| 3300019854 assembled<br>Ga0206370_1116 | 1089 | 0.21 | 1 | k99_59687 | 22 |
| 3300019854 assembled<br>Ga0206370_1117 | 3180 | 0.29 | 2 | k99_65447 | 22 |
| 3300019854 assembled<br>Ga0206370_1118 | 2584 | 0.23 | 3 | k99_65721 | 22 |
| 3300019854 assembled<br>Ga0206370_1119 | 1788 | 0.2 | 3 | k99_71090 | 22 |
| 3300019854 assembled<br>Ga0206370_1120 | 1266 | 0.19 | 1 | k99_80498 | 22 |
| 3300019854 assembled<br>Ga0206370_1121 | 1151 | 0.24 | 2 | k99_92227 | 22 |
| 3300019854 assembled<br>Ga0206370_1122 | 1289 | 0.26 | 1 | k99_96414 | 22 |
| 3300019854 assembled<br>Ga0206370_1123 | 2211 | 0.2 | 5 | k99_100001 | 22 |
| 3300019854 assembled<br>Ga0206370_1124 | 2725 | 0.23 | 2 | k99_101676 | 22 |
| 3300019854 assembled<br>Ga0206370_1125 | 1932 | 0.22 | 4 | k99_109042 | 22 |
| 3300019854 assembled<br>Ga0206370_1126 | 2227 | 0.24 | 1 | k99_118423 | 22 |
| 3300019854 assembled<br>Ga0206370_1127 | 2392 | 0.26 | 5 | k99_119125 | 22 |
| 3300019854 assembled<br>Ga0206370_1128 | 2866 | 0.22 | 8 | k99_130213 | 22 |
| 3300019854 assembled<br>Ga0206370_1129 | 1510 | 0.23 | 1 | k99_138420 | 22 |
| 3300019854 assembled<br>Ga0206370_1130 | 1632 | 0.23 | 2 | k99_141749 | 22 |
| 3300019854 assembled<br>Ga0206370_1131 | 1793 | 0.24 | 1 | k99_143710 | 22 |
| 3300019854 assembled<br>Ga0206370_1132 | 1117 | 0.23 | 1 | k99_150350 | 22 |
| 3300019854 assembled<br>Ga0206370_1133 | 1120 | 0.25 | 2 | k99_162950 | 22 |
| 3300019854 assembled<br>Ga0206370_1134 | 3642 | 0.23 | 5 | k99_182603 | 22 |
| 3300019854 assembled<br>Ga0206370_1135 | 4654 | 0.22 | 7 | k99_189503 | 22 |
| 3300019854 assembled<br>Ga0206370_1136 | 2693 | 0.24 | 4 | k99_195522 | 22 |
| 3300019854 assembled<br>Ga0206370_1137 | 1714 | 0.26 | 1 | k99_196314 | 22 |
| 3300019854 assembled<br>Ga0206370_1138 | 1202 | 0.21 | 2 | k99_199939 | 22 |
| 3300019854 assembled<br>Ga0206370_1139 | 1740 | 0.21 | 5 | k99_205945 | 22 |
| 3300019854 assembled<br>Ga0206370_1140 | 1028 | 0.21 | 4 | k99_217168 | 22 |
| 3300019854 assembled<br>Ga0206370_1141 | 1172 | 0.25 | 1 | k99_231344 | 22 |
| 3300019854 assembled<br>Ga0206370_1142 | 1612 | 0.23 | 2 | k99_234986 | 22 |
| 3300019854 assembled<br>Ga0206370_1143 | 3723 | 0.25 | 3 | k99_238015 | 22 |
| 3300019854 assembled<br>Ga0206370_1144 | 7290 | 0.26 | 9 | k99_238881 | 22 |
| 3300019854 assembled<br>Ga0206370_1145 | 1291 | 0.23 | 2 | k99_264829 | 22 |
| 3300019854 assembled<br>Ga0206370_1146 | 2421 | 0.2 | 3 | k99_295488 | 22 |
| 3300019854 assembled<br>Ga0206370_1147 | 2319 | 0.26 | 4 | k99_296354 | 22 |
| 3300019854 assembled<br>Ga0206370_1148 | 1178 | 0.24 | 3 | k99_298409 | 22 |
| 3300019854 assembled<br>Ga0206370_1149 | 7736 | 0.24 | 14 | k99_311440 | 22 |
| 3300019854 assembled<br>Ga0206370_1150 | 3891 | 0.21 | 6 | k99_311912 | 22 |
| 3300019854 assembled<br>Ga0206370_1151 | 1082 | 0.28 | 2 | k99_312971 | 22 |
| 3300019854 assembled<br>Ga0206370_1152 | 1578 | 0.28 | 1 | k99_325899 | 22 |
| 3300019854 assembled<br>Ga0206370_1153 | 1125 | 0.24 | 3 | k99_332670 | 22 |
| 3300019854 assembled<br>Ga0206370_1154 | 1317 | 0.2 | 1 | k99_333243 | 22 |
| 3300019854 assembled<br>Ga0206370_1155 | 2827 | 0.26 | 3 | k99_345036 | 22 |
| 3300019854 assembled<br>Ga0206370_1156 | 1574 | 0.23 | 3 | k99_369537 | 22 |
| 3300019854 assembled<br>Ga0206370_1157 | 1500 | 0.22 | 5 | k99_373025 | 22 |
| 3300019854 assembled<br>Ga0206370_1158 | 3011 | 0.23 | 8 | k99_382777 | 22 |
| 3300019854 assembled<br>Ga0206370_1159 | 3658 | 0.24 | 7 | k99_401494 | 22 |
| 3300019854 assembled<br>Ga0206370_1160 | 2117 | 0.24 | 2 | k99_402263 | 22 |
| 3300019854 assembled<br>Ga0206370_1161 | 1744 | 0.24 | 1 | k99_402841 | 22 |
| 3300019854 assembled<br>Ga0206370_1162 | 1940 | 0.29 | 2 | k99_405510 | 22 |
| 3300019854 assembled<br>Ga0206370_1163 | 1181 | 0.21 | 1 | k99_408784 | 22 |
| 3300019854 assembled<br>Ga0206370_1164 | 1979 | 0.2 | 5 | k99_416102 | 22 |
| 3300019854 assembled<br>Ga0206370_1165 | 1755 | 0.21 | 3 | k99_426401 | 22 |
| 3300019854 assembled<br>Ga0206370_1166 | 1317 | 0.2 | 3 | k99_428527 | 22 |
| 3300019854 assembled<br>Ga0206370_1167 | 1060 | 0.17 | 2 | k99_428797 | 22 |
| 3300019854 assembled<br>Ga0206370_1168 | 1479 | 0.22 | 2 | k99_436752 | 22 |
| 3300019854 assembled<br>Ga0206370_1169 | 2373 | 0.22 | 7 | k99_444757 | 22 |
| 3300019854 assembled<br>Ga0206370_1170 | 3205 | 0.23 | 8 | k99_447595 | 22 |
| 3300019854 assembled<br>Ga0206370_1171 | 2655 | 0.23 | 3 | k99_462848 | 22 |
| 3300019854 assembled<br>Ga0206370_1172 | 1598 | 0.26 | 2 | k99_463234 | 22 |
| 3300019854 assembled<br>Ga0206370_1173 | 1333 | 0.22 | 5 | k99_471900 | 22 |
| 3300019854 assembled<br>Ga0206370_1174 | 1319 | 0.22 | 2 | k99_479461 | 22 |
| 3300019854 assembled<br>Ga0206370_1175 | 55827 | 0.39 | 78 | k99_264868 | 3 |
| 3300019854 assembled<br>Ga0206370_1176 | 31696 | 0.35 | 50 | k99_449216 | 13 |
| 3300019854 assembled<br>Ga0206370_1177 | 43776 | 0.38 | 65 | k99_302609 | 19 |
| 3300019854 assembled<br>Ga0206370_1178 | 24175 | 0.45 | 40 | k99_324565 | 5 |
| 3300019854 assembled<br>Ga0206370_1179 | 120359 | 0.56 | 136 | k99_196508 | 17 |

**Supplemental Table 2:** Presence of CRISPR spacer matches from IMG/VR spacer database to oral phage.

| Contig | Oral Phage | Genome accession of spacer | Species |
| --- | --- | --- | --- |
| k99_345784 | 7 | AEPG01000403 | Neisseria sicca DS1 |
| k99_345784 | 7 | NZ_AFAY01000002 | Neisseria bacilliformis ATCC BAA-1200 |
| k99_345784 | 7 | NZ_AEWV01000008 | Kingella denitrificans ATCC 33394 |
| k99_347722 | 8 | Ga0077313_161 | Haemophilus haemolyticus 1P26 |
| k99_283470 | 16 | AWSF01000235 | Actinomyces sp. oral taxon 172 str. F0311 |
| k99_283470 | 16 | ALIY01000071 | Actinomyces sp. ICM58 |
| k99_66521 | 16 | NZ_AAYI02000004 | Actinomyces odontolyticus ATCC 17982 |
| k99_313503 | 18 | NZ_ACYT02000015 | Actinomyces odontolyticus F0309 |
| k99_368282 | 18 | ALCA01000063 | Actinomyces sp. ICM47 |
| k99_302609 | 19 | ALOA01000009 | Lachnoanaerobaculum sp. OBRC5-5 |
| k99_302609 | 19 | AJGH01000032 | Lachnoanaerobaculum saburreum F0468 |
| k99_302609 | 19 | NZ_AGRL01000181 | Lachnospiraceae bacterium oral taxon 082 str. F0431 |
| k99_302609 | 19 | NZ_AGRL01000179 | Lachnospiraceae bacterium oral taxon 082 str. F0431 |
| k99_302609 | 19 | NZ_AGRL01000179 | Lachnospiraceae bacterium oral taxon 082 str. F0431 |
| k99_302609 | 19 | AJGH01000032 | Lachnoanaerobaculum saburreum F0468 |
| k99_170084 | 21 | AFFN01000027 | Streptococcus sanguinis SK355 |
| k99_249821 | 24 | AQTU01000003 | Streptococcus mitis 13/39 |
| k99_293096 | 24 | ATAA01000014 | Streptococcus mitis 18/56 |
| k99_79926 | 24 | NZ_GL732467 | Streptococcus sp. C300 |
| k99_79926 | 24 | NZ_GL732467 | Streptococcus sp. C300 |
| k99_79926 | 24 | JPFT01000005 | Streptococcus mitis |
| k99_79926 | 24 | ATAA01000014 | Streptococcus mitis 18/56 |
| k99_79926 | 24 | ALST01000003 | Streptococcus agalactiae GB00084 |
| k99_122647 | 25 | JPFT01000005 | Streptococcus mitis |
| k99_82015 | 25 | AJRG02000005 | Streptococcus sp. GMD5S |
| k99_82015 | 25 | AJRH01000303 | Streptococcus sp. GMD6S |
| k99_82015 | 25 | AJRA01000599 | Streptococcus sp. GMD2S |
| k99_82015 | 25 | AJQY01000467 | Streptococcus sp. GMD1S |

**Supplemental Table 3:** Similarities of oral phage to bacterial isolate genomes.

| Phage number | Best Hit Accession(s) | Strain | Percent Coverage | Percent Identity |
| --- | --- | --- | --- | --- |
| 1 | NZ_JH414504.1 | Oribacterium asaccharolyticum ACB7 | 84.2% | 79.1% |
| 4 | NZ_LZMW01000016.1 | Haemophilus parainfluenzae strain CCUG 58848 | 100.0% | 100.0% |
|  | NZ_JUTJ01000008.1 | Haemophilus parainfluenzae strain 777_HPAR | 100.0% | 100.0% |
|  | NZ_JWCE01000055.1 | Haemophilus parainfluenzae strain 1128_HPAR | 100.0% | 100.0% |
|  | NZ_JUTE01000022.1 | Haemophilus influenzae strain 781_HINF | 100.0% | 100.0% |
|  | NZ_KV820357.1 | Haemophilus sp. HMSC068C11 | 100.0% | 100.0% |
|  | NZ_LZMX01000030.1 | Haemophilus parainfluenzae strain CCUG 62654 | 100.0% | 99.9% |
| 5 | NZ_MUXX01000020.1 | Haemophilus paraphrohaemolyticus strain CCUG 3718 | 85.2% | 91.9% |
| 6 | NZ_KQ959948.1 | Gemella haemolysans strain DNF01167 | 63.5% | 89.1% |
|  | NZ_KQ959944.1 |  |  |  |
| 7 | NZ_FFBA01000005.1 | Neisseria meningitidis strain 2842STDY5881411 | 87.2% | 95.1% |
| 8 | NZ_LVZF01000011.1 | Haemophilus influenzae strain GE49 | 80.0% | 91.7% |
| 13 | NZ_CRPU01000013.1 | Streptococcus pneumoniae strain SMRU824 | 57.7% | 72.2% |
| 16 | NZ_JDFI01000000 | Actinomyces sp. ICM54 | 94.3% | 91.1% |
| 23 | NZ_FEBQ01000017.1 | Streptococcus pneumoniae strain 2842STDY5643923 | 61.3% | 72.7% |
|  | NZ_NCVF01000020.1 | Streptococcus mitis strain RH_50275_09 |  |  |
|  | NZ_KV831007.1 | Streptococcus sp. HMSC063B03 |  |  |

**Supplemental Table 4:** Similarities of ends of phage contigs to bacterial isolate genomes.

| Phage number | Phage Contig | Bacterial Contig | Bacterial Strain | Phage nt range | Bacterial nt range | % identity |
| --- | --- | --- | --- | --- | --- | --- |
| 12 | k99_417291 | NZ_AFUJ01000001.1 | *Veillonella* sp. oral taxon 780 | 1-175 | 17,280-17,454 | 94% |
|  |  |  |  | 38,845-39,582 | 17,442-18,179 | 93% |
| 12 | k99_417291 | NZ_KQ960771.1 | *Veillonella* sp. DNF00869 | 1-175 | 167,420-167,594 (complement) | 97% |
|  |  |  |  | 38,845-39,582 | 166,695-167,432 (complement) | 92% |
| 19 | k99_302609 | NZ_ALJL01000036.1 | *Lachnoanaerobaculum* sp. ICM7 | 43-1694 | 186,777-188,427 (complement) | 99% |
|  |  |  |  | 43,683-43,776 | 188,339-188,432 (complement) | 95% |

**Supplemental Table 5:** Integrase and related genes.

| Oral Phage Number | Contig | IMG gene number | Description |
| --- | --- | --- | --- |
| 5 | 324565 | 117829 | Integrase core domain-containing protein |
| 6 | 326664 | 102835 | Integrase |
| 7 | 345784 | 110540 | Integrase core domain-containing protein |
| 8 | 347722 | 10162 | Integrase |
| 12 | 417291 | 111041 | Integrase |
| 16 | 66521 | 110713 | Integrase |
| 17 | 196508 | 1179104 | Predicted site-specific integrase-resolvase |
| 18 | 368282 | 101535 | Holliday junction resolvase RusA (prophage-encoded endonuclease) |
| 19 | 302609 | 117750 | Integrase |
|  |  | 117760 | Integrase |
|  |  | 11771 | Transposase |
|  |  | 117764 | site-specific DNA recombinase |
|  |  | 117743 | Holliday junction resolvase RusA (prophage-encoded endonuclease) |
|  |  | 117740 | Site-specific recombinase XerD |
|  |  | 117739 | Site-specific recombinase XerD |
| 20 | 281895 | 10903 | Holliday junction resolvase RuvABC<br>endonuclease subunit |
| 22 | 15350 | 11321 | putative transposase |
| 22 | 311440 | 114914 | putative transposase |
| 22 | 401494 | 11595 | putative transposase |

**Supplemental Table 6:** Viral Cluster Analysis.

| Oral Phage | Viral Clusters | Singletons |
| --- | --- | --- |
| Phage 1 | vc_17442 |  |
| Phage 2 |  |  |
| Phage 3 | vc_17051 |  |
| Phage 4 | vc_17090 |  |
| Phage 5 | vc_19240 | sg_108845 |
| Phage 6 |  |  |
| Phage 7 | vc_17282 |  |
| Phage 8 | vc_17996 |  |
| Phage 9 |  |  |
| Phage 10 |  | sg_106662, sg_106665 |
| Phage 11 |  |  |
| Phage 12 | vc_18231 |  |
| Phage 13 |  |  |
| Phage 14 | vc_18188 | sg_106653, sg_113186 |
| Phage 15 | vc_19538 |  |
| Phage 16 |  | sg_107067, sg_112965 |
| Phage 17 | vc_17438 | sg_106290 |
| Phage 18 | vc_18685 |  |
| Phage 19 |  |  |
| Phage 20 | vc_17030, vc_19658 | sg_103863, sg_104523, sg_111201, sg_111213, sg_112205, sg_112825, sg_112865, sg_113057 |
| Phage 21 | vc_18971 | sg_106366 |
| Phage 22 | vc_19325, vc_19326, vc_19328,<br>vc_19332 | sg_104258, sg_104266, sg_104271, sg_106739,<br>sg_104701, sg_106731, sg_107388, sg_108349,<br>sg_112716 |
| Phage 23 |  |  |
| Phage 24 | vc_19392 | sg_104304, sg_107838, sg_108058, sg_111075,<br>sg_113083 |
| Phage 25 | vc_19392 | sg_104304, sg_107838, sg_108058, sg_111075,<br>sg_113083 |

Note: viral cluster numbers are specific to version IMG\_VR\_2017-01-01\_2 of IMG/VR available at: https://genome.jgi.doe.gov/portal/IMG_VR/IMG_VR.download.html

## REFERENCES

1. Tanner ACR, Kressirer CA, Faller LL. 2016. Understanding Caries From the Oral Microbiome Perspective. J Calif Dent Assoc 44:437–446.

2. Griffen, Ann, L, Beall, Clifford, J, Campbell, James, H, Firestone, Noah, D, Kumar, Purnima, S, Yang, Zamin, K, Podar M, Leys, Eugene J. 2012. Distinct and complex bacterial profiles in human periodontitis and health revealed by 16S pyrosequencing. ISME J 6:1176–1185.

3. Abusleme L, Dupuy AK, Dutzan N, Silva N, Burleson JA, Strausbaugh LD, Gamonal J, Diaz PI. 2013. The subgingival microbiome in health and periodontitis and its relationship with community biomass and inflammation. ISME J 7:1016–1025.

4. Szafrański SP, Winkel A, Stiesch M. 2017. The use of bacteriophages to biocontrol oral biofilms. J Biotechnol 250:29–44.

5. Wade WG. 2013. The oral microbiome in health and disease. Pharmacol Res 69:137–143.

6. Edlund A, Santiago-Rodriguez TM, Boehm TK, Pride DT. 2015. Bacteriophage and their potential roles in the human oral cavity. J Oral Microbiol 7:27423.

7. Abeles SR, Robles-Sikisaka R, Ly M, Lum AG, Salzman J, Boehm TK, Pride DT. 2014. Human oral viruses are personal, persistent and gender-consistent. ISME J 8:1753.

8. Ly M, Abeles SR, Boehm TK, Robles-Sikisaka R, Naidu M, Santiago-Rodriguez T, Pride DT. 2014. Altered oral viral ecology in association with periodontal disease. MBio 5:e01133–14.

9. Pride DT, Salzman J, Haynes M, Rohwer F, Davis-Long C, White RA 3rd, Loomer P, Armitage GC, Relman DA. 2012. Evidence of a robust resident bacteriophage population revealed through analysis of the human salivary virome. ISME J 6:915–926.

10. Abeles SR, Ly M, Santiago-Rodriguez TM, Pride DT. 2015. Effects of Long Term Antibiotic Therapy on Human Oral and Fecal Viromes. PLoS One 10:e0134941.

11. Pride DT, Salzman J, Relman DA. 2012. Comparisons of clustered regularly interspaced short palindromic repeats and viromes in human saliva reveal bacterial adaptations to salivary viruses. Environ Microbiol 14:2564–2576.

12. Robles-Sikisaka R, Naidu M, Ly M, Salzman J, Abeles SR, Boehm TK, Pride DT. 2014. Conservation of streptococcal CRISPRs on human skin and saliva. BMC Microbiol 14:146.

13. Roux S, Enault F, Hurwitz BL, Sullivan MB. 2015. VirSorter: mining viral signal from microbial genomic data. PeerJ 3:e985.

14. Paez-Espino D, Pavlopoulos GA, Ivanova NN, Kyrpides NC. 2017. Nontargeted virus sequence discovery pipeline and virus clustering for metagenomic data. Nat Protoc 12:1673–1682.

15. Paez-Espino D, Eloe-Fadrosh EA, Pavlopoulos GA, Thomas AD, Huntemann M, Mikhailova N, Rubin E, Ivanova NN, Kyrpides NC. 2016. Uncovering Earth’s virome. Nature 536:425–430.

16. Alneberg J, Bjarnason BS, de Bruijn I, Schirmer M, Quick J, Ijaz UZ, Lahti L, Loman NJ, Andersson AF, Quince C. 2014. Binning metagenomic contigs by coverage and composition. Nat Methods 11:1144–1146.

17. Truong DT, Franzosa EA, Tickle TL, Scholz M, Weingart G, Pasolli E, Tett A, Huttenhower C, Segata N. 2015. MetaPhlAn2 for enhanced metagenomic taxonomic profiling. Nat Methods 12:902–903.

18. Bolger AM, Lohse M, Usadel B. 2014. Trimmomatic: a flexible trimmer for Illumina sequence data. Bioinformatics 30:2114–2120.

19. Li H. 2013. Aligning sequence reads, clone sequences and assembly contigs with BWA-MEM. arXiv [q-bioGN].

20. Li D, Liu C-M, Luo R, Sadakane K, Lam T-W. 2015. MEGAHIT: an ultra-fast single-node solution for large and complex metagenomics assembly via succinct de Bruijn graph. Bioinformatics 31:1674–1676.

21. Mikheenko A, Saveliev V, Gurevich A. 2016. MetaQUAST: evaluation of metagenome assemblies. Bioinformatics 32:1088–1090.

22. Buchfink B, Xie C, Huson DH. 2015. Fast and sensitive protein alignment using DIAMOND. Nat Methods 12:59–60.

23. Huson DH, Beier S, Flade I, Górska A, El-Hadidi M, Mitra S, Ruscheweyh H-J, Tappu R. 2016. MEGAN Community Edition - Interactive Exploration and Analysis of Large-Scale Microbiome Sequencing Data. PLoS Comput Biol 12:e1004957.

24. Li H, Handsaker B, Wysoker A, Fennell T, Ruan J, Homer N, Marth G, Abecasis G, Durbin R, 1000 Genome Project Data Processing Subgroup. 2009. The Sequence Alignment/Map format and SAMtools. Bioinformatics 25:2078–2079.

25. Cock PJ, Antao T, Chang JT, Chapman BA, Cox CJ, Dalke A, Friedberg I, Hamelryck T, Kauff F, Wilczynski B, de Hoon MJ. 2009. Biopython: freely available Python tools for computational molecular biology and bioinformatics. Bioinformatics 25:1422–1423.

26. Garrison E, Marth G. 2012. Haplotype-based variant detection from short-read sequencing. arXiv [q-bioGN].

27. Schloissnig S, Arumugam M, Sunagawa S, Mitreva M, Tap J, Zhu A, Waller A, Mende DR, Kultima JR, Martin J, Kota K, Sunyaev SR, Weinstock GM, Bork P. 2013. Genomic variation landscape of the human gut microbiome. Nature 493:45–50.

28. Chen I-MA, Markowitz VM, Chu K, Palaniappan K, Szeto E, Pillay M, Ratner A, Huang J, Andersen E, Huntemann M, Varghese N, Hadjithomas M, Tennessen K, Nielsen T, Ivanova NN, Kyrpides NC. 2017. IMG/M: integrated genome and metagenome comparative data analysis system. Nucleic Acids Res 45:D507–D516.

29. Bolduc B, Jang HB, Doulcier G, You Z-Q, Roux S, Sullivan MB. 2017. vConTACT: an iVirus tool to classify double-stranded DNA viruses that infect Archaea and Bacteria. PeerJ 5:e3243.

30. Lowe TM, Eddy SR. 1997. tRNAscan-SE: a program for improved detection of transfer RNA genes in genomic sequence. Nucleic Acids Res 25:955–964.

31. Camacho C, Coulouris G, Avagyan V, Ma N, Papadopoulos J, Bealer K, Madden TL. 2009. BLAST+: architecture and applications. BMC Bioinformatics 10:421.

32. Paez-Espino D, Chen I-MA, Palaniappan K, Ratner A, Chu K, Szeto E, Pillay M, Huang J, Markowitz VM, Nielsen T, Huntemann M, K Reddy TB, Pavlopoulos GA, Sullivan MB, Campbell BJ, Chen F, McMahon K, Hallam SJ, Denef V, Cavicchioli R, Caffrey SM, Streit WR, Webster J, Handley KM, Salekdeh GH, Tsesmetzis N, Setubal JC, Pope PB, Liu W-T, Rivers AR, Ivanova NN, Kyrpides NC. 2017. IMG/VR: a database of cultured and uncultured DNA Viruses and retroviruses. Nucleic Acids Res 45:D457–D465.

33. Skennerton CT, Imelfort M, Tyson GW. 2013. Crass: identification and reconstruction of CRISPR from unassembled metagenomic data. Nucleic Acids Res 41:e105.

34. Langmead B, Salzberg SL. 2012. Fast gapped-read alignment with Bowtie 2. Nat Methods 9:357–359.

35. Delisle AL, Barcak GJ, Guo M. 2006. Isolation and expression of the lysis genes of Actinomyces naeslundii phage Av-1. Appl Environ Microbiol 72:1110–1117.

36. Martín AC, López R, García P. 1996. Analysis of the complete nucleotide sequence and functional organization of the genome of Streptococcus pneumoniae bacteriophage Cp-1. J Virol 70:3678–3687.

37. Romero P, López R, García E. 2004. Genomic organization and molecular analysis of the inducible prophage EJ-1, a mosaic myovirus from an atypical pneumococcus. Virology 322:239–252.

38. Oh J, Byrd AL, Park M, Program NCS, Kong HH, Segre JA. 2016. Temporal Stability of the Human Skin Microbiome. Cell 165:854–866.

39. Momeni SS, Whiddon J, Cheon K, Ghazal T, Moser SA, Childers NK. 2016. Genetic Diversity and Evidence for Transmission of Streptococcus mutans by DiversiLab rep-PCR. J Microbiol Methods 128:108–117.

40. Beall CJ, Campbell AG, Dayeh DM, Griffen AL, Podar M, Leys EJ. 2014. Single Cell Genomics of Uncultured, Health-Associated *Tannerella BU063* (Oral Taxon 286) and Comparison to the Closely Related Pathogen *Tannerella forsythia*. PLoS One 9:e89398.

41. Beall CJ, Campbell AG, Griffen AL, Podar M, Leys EJ. 2018. Genomics of the Uncultivated, Periodontitis-Associated Bacterium Tannerella sp. BU045 (Oral Taxon 808). mSystems 3.

42. Šimoliūnas E, Kaliniene L, Stasilo M, Truncaitė L, Zajančkauskaitė A, Staniulis J, Nainys J, Kaupinis A, Valius M, Meškys R. 2014. Isolation and characterization of vB_ArS-ArV2 - first Arthrobacter sp. infecting bacteriophage with completely sequenced genome. PLoS One 9:e111230.

43. Redmond WB, Cater JC. 1960. A bacteriophage specific for Mycobacterium tuberculosis, varieties hominis and bovis. Am Rev Respir Dis 82:781–786.

44. Pope WH, Bowman CA, Russell DA, Jacobs-Sera D, Asai DJ, Cresawn SG, Jacobs WR, Hendrix RW, Lawrence JG, Hatfull GF, Science Education Alliance Phage Hunters Advancing Genomics and Evolutionary Science, Phage Hunters Integrating Research and Education, Mycobacterial Genetics Course. 2015. Whole genome comparison of a large collection of mycobacteriophages reveals a continuum of phage genetic diversity. Elife 4:e06416.

45. Meijer WJ, Horcajadas JA, Salas M. 2001. Phi29 family of phages. Microbiol Mol Biol Rev 65:261–87; second page, table of contents.

46. Ronda C, López R, García E. 1981. Isolation and characterization of a new bacteriophage, Cp-1, infecting Streptococcus pneumoniae. J Virol 40:551–559.

47. Yuan Y, Gao M. 2017. Jumbo Bacteriophages: An Overview. Front Microbiol 8:403.

48. Timms AR, Cambray-Young J, Scott AE, Petty NK, Connerton PL, Clarke L, Seeger K, Quail M, Cummings N, Maskell DJ, Thomson NR, Connerton IF. 2010. Evidence for a lineage of virulent bacteriophages that target Campylobacter. BMC Genomics 11:214.

49. Kono N, Arakawa K, Tomita M. 2011. Comprehensive prediction of chromosome dimer resolution sites in bacterial genomes. BMC Genomics 12:19.

50. Askora A, Abdel-Haliem MEF, Yamada T. 2012. Site-specific recombination systems in filamentous phages. Mol Genet Genomics 287:525–530.

51. Albertsen M, Hugenholtz P, Skarshewski A, Nielsen KL, Tyson GW, Nielsen PH. 2013. Genome sequences of rare, uncultured bacteria obtained by differential coverage binning of multiple metagenomes. Nat Biotechnol 31:533–538.

52. Duranti S, Lugli GA, Mancabelli L, Armanini F, Turroni F, James K, Ferretti P, Gorfer V, Ferrario C, Milani C, Mangifesta M, Anzalone R, Zolfo M, Viappiani A, Pasolli E, Bariletti I, Canto R, Clementi R, Cologna M, Crifò T, Cusumano G, Fedi S, Gottardi S, Innamorati C, Masè C, Postai D, Savoi D, Soffiati M, Tateo S, Pedrotti A, Segata N, van Sinderen D, Ventura M. 2017. Maternal inheritance of bifidobacterial communities and bifidophages in infants through vertical transmission. Microbiome 5:66.

53. Ly M, Jones MB, Abeles SR, Santiago-Rodriguez TM, Gao J, Chan IC, Ghose C, Pride DT. 2016. Transmission of viruses via our microbiomes. Microbiome 4:64.

54. Milani C, Casey E, Lugli GA, Moore R, Kaczorowska J, Feehily C, Mangifesta M, Mancabelli L, Duranti S, Turroni F, Bottacini F, Mahony J, Cotter PD, McAuliffe FM, van Sinderen D, Ventura M. 2018. Tracing mother-infant transmission of bacteriophages by means of a novel analytical tool for shotgun metagenomic datasets: METAnnotatorX. Microbiome 6:145.

